# Functional potential and evolutionary response to long-term heat selection of bacterial associates of coral photosymbionts

**DOI:** 10.1101/2023.08.04.552070

**Authors:** Justin Maire, Gayle K. Philip, Jadzia Livingston, Louise M. Judd, Linda L. Blackall, Madeleine J. H. van Oppen

## Abstract

Corals rely on a wide range of microorganisms for their functioning, including intracellular dinoflagellates (Symbiodiniaceae) and bacteria. Marine heatwaves trigger the loss of Symbiodiniaceae from coral tissues - coral bleaching - often leading to death. While coral-bacteria interactions are widely studied, Symbiodiniaceae-bacteria interactions have remained largely uninvestigated. Here, we provide a genomic analysis of 49 bacteria, spanning 16 genera, that closely associate with six cultured Symbiodiniaceae species. We analyzed bacterial functional potential by focusing on potentially beneficial functions for the Symbiodiniaceae host, including B vitamin synthesis and antioxidant abilities, which may be crucial for Symbiodiniaceae heat tolerance and in turn coral resistance to thermal bleaching. These analyses suggest a wide potential for B vitamin synthesis and the scavenging of reactive oxygen species (through the production of carotenoids or antioxidant enzymes), and possibly the transfer of organic carbon to host cells. Single nucleotide polymorphism analysis between bacteria isolated from wild-type and heat-evolved Symbiodiniaceae cultures revealed that exposure to long-term elevated temperature has resulted in mutations in genes known to be involved in host-symbiont interactions, such as secretion systems. Climate change may therefore modify how Symbiodiniaceae and bacteria interact. This study provides an overview of the possible roles of Symbiodiniaceae-associated bacteria in Symbiodiniaceae functioning and heat tolerance, reinforcing the need for further studies of such interactions to fully understand coral biology and climate resilience.

**Importance:** Symbiotic microorganisms are crucial for the survival of corals and their resistance to coral bleaching in the face of climate change. However, the impact of microbe-microbe interactions on coral functioning is mostly unknown, but could be essential factors for coral adaption to future climates. Here, we investigated interactions between cultured dinoflagellates of the Symbiodiniaceae family, essential photosymbionts of corals, and associated bacteria. By assessing the genomic potential of 49 bacteria, we found that they are likely beneficial for Symbiodiniaceae, through the production of B vitamins and antioxidants. Additionally, bacterial genes involved in host-symbiont interactions, such as secretion systems, accumulated mutations following long-term exposure to heat, suggesting symbiotic interactions may change under climate change. This highlights the importance of microbe-microbe interactions in coral functioning.

## Introduction

Coral reefs are valuable ecosystems, as they provide a home to about a third of all eukaryotic marine species as well as benefits to human populations, such as food security and coastal protection (1, 2). Reef-building corals rely on a diverse suite of microorganisms for their survival (3–5). Endosymbiotic dinoflagellates of the Symbiodiniaceae family (Suessiales, Dinophyta) are probably the most crucial and most studied member of this microbial consortium (6). By providing photosynthate to their host, Symbiodiniaceae meet most of the energy needs of corals (7), and are essential for the health of coral reefs. Unfortunately, this symbiotic association is extremely fragile and is particularly sensitive to temperature fluctuations. Increases in sea surface temperatures can result in the loss of Symbiodiniaceae from coral tissues, a process known as coral bleaching, which can result in coral death (8). The main mechanism thought to trigger coral bleaching is an increase in the production of reactive oxygen species (ROS) by Symbiodiniaceae, which leak into coral cells, overwhelm the coral’s antioxidant response, and trigger a cellular cascade that results in Symbiodiniaceae loss (9). Mass coral bleaching events are increasing in frequency and intensity because of anthropogenic climate change (10), thus highlighting the need to increase knowledge on coral and Symbiodiniaceae biology and to develop novel coral conservation and restoration methods, while greenhouse gas emissions are curbed.

Corals also house a wide diversity of bacteria, which colonize all microhabitats within their host (4, 11–13). Putative bacterial functions include nutrient cycling and protection against pathogens (4, 12). Recently, Symbiodiniaceae were also found to associate with bacteria, both extra- and intracellularly (14–16). Symbiodiniaceae-associated bacterial communities are highly diverse and change after both short-term thermal stress (17–19) and long-term heat selection (19), pointing at a potential bacterial involvement in Symbiodiniaceae thermal tolerance. Two studies have shown that inoculation of Symbiodiniaceae cultures with a single bacterial strain (belonging to the *Muricauda* and *Roseovarius* genera, respectively) can enhance Symbiodiniaceae (*Durusdinium* sp. and *Breviolum minutum*, respectively) performance under temperature and/or light stress (18, 20). In the case of *Durusdinium/Muricauda*, this was attributed to the production of a carotenoid by *Muricauda*, a pigment able to scavenge ROS (20). Nevertheless, the functions and functional potential of most Symbiodiniaceae-associated bacteria remain mostly unexplored (13, 21).

Here, we provide the first comparative genomic analysis of Symbiodiniaceae-associated bacteria. From a culture collection of more than 200 bacterial isolates, cultured from six Symbiodiniaceae species, we sequenced the genomes of 49 representative isolates. We present an in-depth analysis of bacterial functional potential that may be beneficial for Symbiodiniaceae hosts, such as B vitamin synthesis and ROS scavenging. We also analyze the effect of Symbiodiniaceae thermal selection on bacterial genomic evolution. This study provides an important step to understanding the contribution of bacteria to Symbiodiniaceae functioning, and in turn their roles in coral health and climate resilience.

## Results and discussion

### Isolation of bacteria from Symbiodiniaceae cultures

In a previous study, 141 pure bacterial isolates, spanning 20 genera, were obtained from six Symbiodiniaceae species (15). We recently showed that long-term heat selection of the species *Cladocopium proliferum* affected the composition of Symbiodiniaceae-associated bacteria (19), both at the extracellular and intracellular level (22). Thus, we expanded our culture collection by pure culturing bacteria from one wild-type (WT10) and two heat-evolved (SS03 and SS08) *C. proliferum* strains. For each *C. proliferum* strain, samples were washed, and either bead-beat, in order to release potentially intracellular bacteria, or not. A total of 65 pure bacterial isolates were obtained and identified through 16S rRNA gene sequence analysis (Table S1). These newly obtained isolates spanned ten genera, five of which were previously isolated from Symbiodiniaceae cultures (15). Four genera (*Marinobacter, Muricauda, Mamelliela,* and *Roseitalea*) were cultured from all three *C. proliferum* strains.

Six genera (*Janibacter, Aestuariicoccus, Muricauda, Roseibium*, *Roseitalea*, and *Oceaniradius*) were isolated exclusively from *C. proliferum* samples that were bead beaten prior to spreading on agar plates, suggesting they may be intracellular associates. Nonetheless, *Muricauda, Roseitalea*, and *Roseibium* (previously referred to as *Labrenzia*) were previously isolated from *C. proliferum* WT10 samples that were not bead beaten (15), suggesting they can be extracellular. Therefore, only *Janibacter, Aestuariicoccus,* and *Oceaniradius* are strong candidates for being intracellular symbionts of *C. proliferum*, while all others are likely extracellular, though still closely associated.

### Genome sequencing of 49 Symbiodiniaceae-associated bacteria

To explore the genomic abilities of bacteria that closely associate with Symbiodiniaceae, the whole genome of 49 isolates were sequenced, assembled, and annotated. The 49 isolates were chosen to span both the bacterial and Symbiodiniaceae diversity of our culture collection. All 49 assembled genomes exhibited a completeness above 98% and contamination below 2.5%, and ranged from 2.98-6.88 Mb in size (Table S2). The taxonomy of these genomes was confirmed with the GTDB-Tk tool, the construction of a phylogenetic tree, and the calculation of average nucleotide and amino acid identity (ANI and AAI, respectively) (Figures 1 and S1, Table S2). This corroborated the taxonomy previously obtained through 16S rRNA gene sequence analysis (see “Taxonomy notes” in Table S2). It is worth noting that, within one genus, isolates from distinct Symbiodiniaceae hosts often show >99.9% ANI and 100% AAI with each other (*e.g.*, *Muricauda*, *Mameliella, Roseitalea*, and *Marinobacter*; Figure S1), suggesting a lack of host-symbiont specificity.

**Figure 1:**
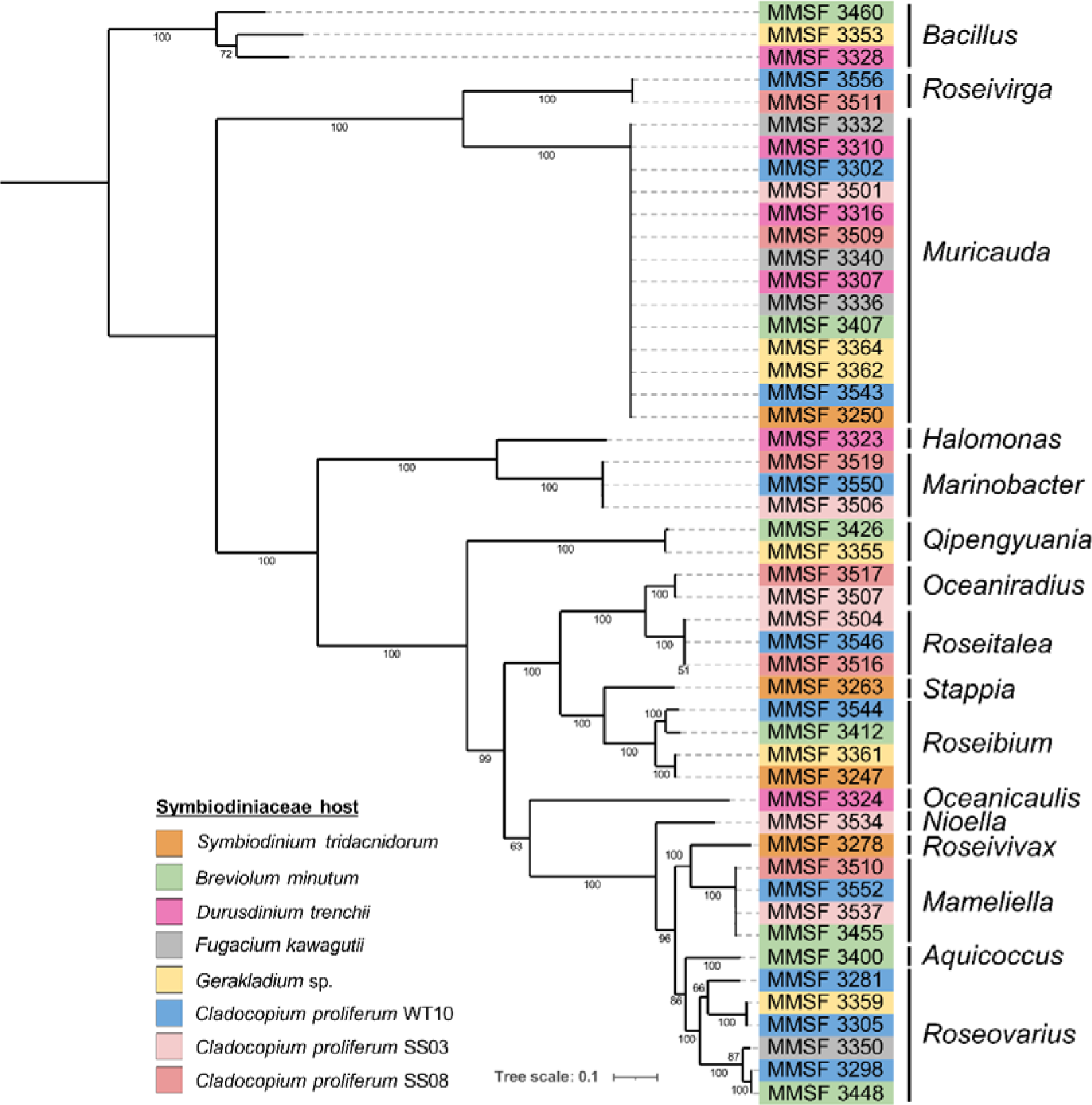
Maximum-likelihood phylogenetic tree of the 49 pure-cultured Symbiodiniaceae-associated bacteria based on 120 marker genes (GTDB-Tk). The 49 isolates were assigned to 16 bacterial genera, as stipulated on the right-hand side. Bootstrap support values based on 1000 replications are provided.

The metabolic abilities of the 49 isolates were reconstructed using a KEGG annotation (Table S3), and pathways of interest are summarized in Figure 2. Pathways that were more than 75% complete were considered functional. Again, no host specificity or co-diversification was observed: isolates of a given genus all had almost identical metabolic potentials, regardless of the Symbiodiniaceae host they were cultured from. This is intriguing as the range of Symbiodiniaceae strains used in the study were originally isolated from four coral species, one sea anemone species, and one giant clam species. This is consistent with the fact that community composition of intracellular bacteria was highly similar across these Symbiodiniaceae cultures (15). Therefore, these bacteria may be opportunistic associates (as opposed to long-term symbionts), perhaps selected during the initial isolation of Symbiodiniaceae or accidentally introduced in Symbiodiniaceae cultures later on, which have been maintained over the years, presumably because they were not detrimental, and potentially beneficial, to their Symbiodiniaceae host.

**Figure 2:**
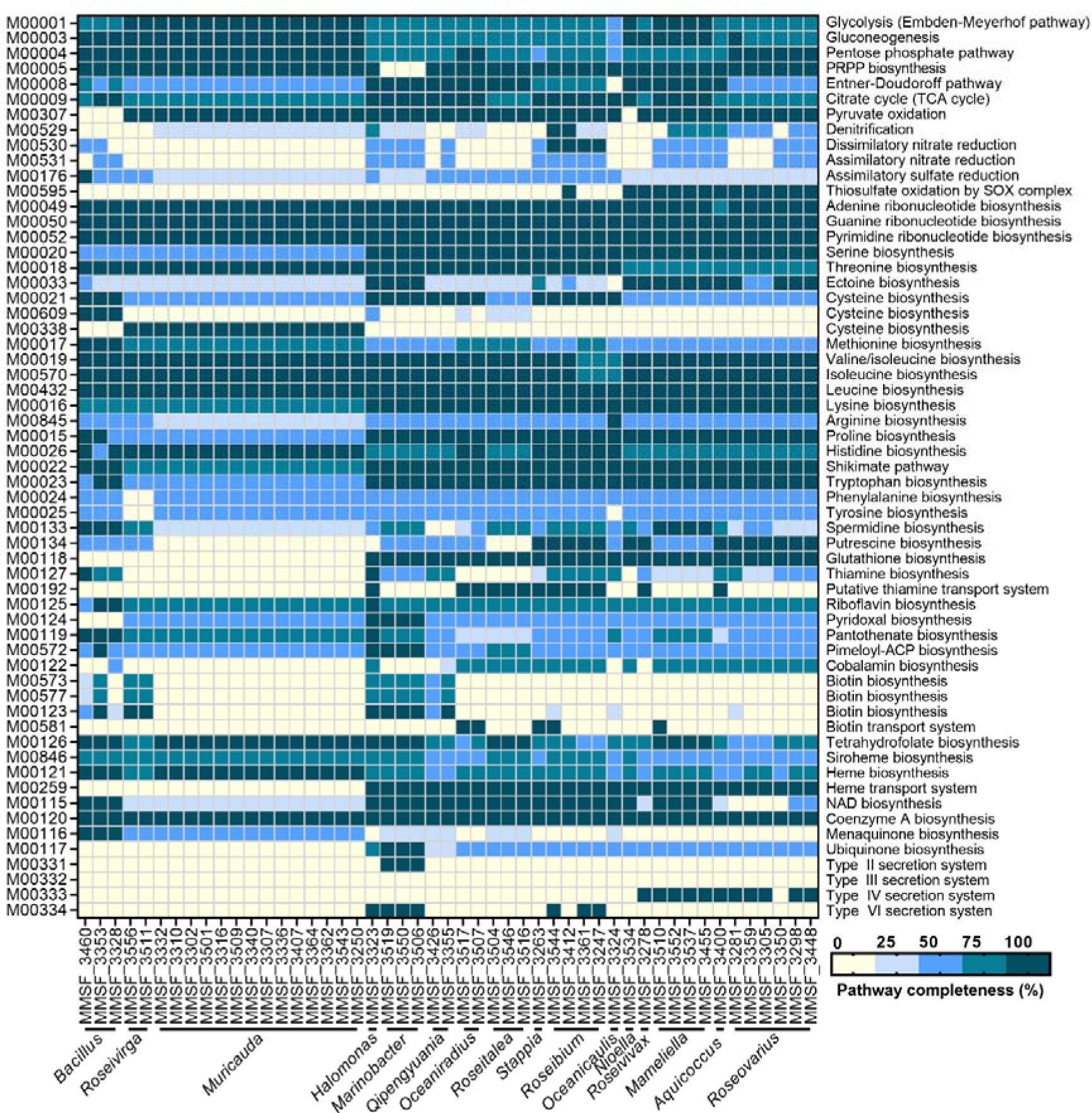
Metabolic potential of 49 Symbiodiniaceae-associated bacteria. The completeness of metabolic pathways (KEGG module database) of interest were annotated using METABOLIC-G and pathway completeness (in %) was estimated in EnrichM. See Table S3 for the full list of pathways.

Almost all isolates possessed all genes for glycolysis, the tricarboxylic acid (TCA) cycle, the pentose phosphate pathway, the biosynthesis of all nucleotides, and the biosynthesis of several amino acids (threonine, valine, isoleucine, leucine, lysine histidine). Only two *Roseibium* isolates had the genes to perform denitrification, while all four *Roseibium* isolates had the genetic potential for dissimilatory nitrate reduction. Denitrifying potential is commonly observed in other *Roseibium* strains (sometimes referred to as *Labrenzia*) (23, 24). Several isolates also possessed complete secretion systems, which may assist in host-bacteria and bacteria-bacteria interactions, including Symbiodiniaceae infection: a type II secretion system (T2SS) was found in three isolates (belonging to *Marinobacter*); a type IV secretion system (T4SS) was found in 11 isolates (belonging to *Roseovarius, Roseivivax, Mameliella, Aquicoccus*); a type VI secretion system (T6SS) was found in seven isolates (belonging to *Halomonas, Marinobacter, Roseibium*). We also interrogated the presence of eukaryotic-like proteins, which may facilitate host-bacteria interactions by mediating bacterial protein-eukaryotic host protein interactions (25), but did not detect any in any of the 49 genomes reported here.

### Potential for B vitamin synthesis

B vitamins are required by most microalgae (26–28), including Symbiodiniaceae (21, 29), and promote algal growth. For example, biotin (vitamin B_7_) is essential for fatty acid metabolism, while thiamine (vitamin B_1_) is important for carbohydrate and amino acid metabolism (27). Commonly used Symbiodiniaceae culture media, such as Daigo’s IMK medium used in this study, contain biotin, thiamine and cobalamin (vitamin B_12_), although some B vitamins have been shown to be provided by bacteria in some cases. For example, *Halomonas* sp. and *Sinorhizobium meliloti* provide cobalamin to the unicellular algae *Porphyridium purpureum* and *Chlamydomonas reinhardtii*, respectively (26, 30). Therefore, we interrogated the completeness of B vitamin biosynthesis pathways in Symbiodiniaceae-associated bacteria (Figure 2, Tables S3 and S4).

All but one isolate were found to be able to synthesize riboflavin (vitamin B_2_), while 43 possessed the genes for tetrahydrofolate (vitamin B_9_) synthesis (only missing in one *Oceaniradius*, two *Roseibium* and three *Roseovarius*), 28 possessed the genes for pantothenate (vitamin B_5_) synthesis (belonging to *Bacillus, Roseivirga, Muricauda, Halomonas, Marinobacter, Oceanicaulis, Mameliella*) and 23 isolates showed potential to synthesize cobalamin (belonging to *Halomonas, Oceaniradius, Roseitalea, Stappia, Roseovarius, Mameliella, Aquicoccus, Roseibium, Nioella*). The potential for the synthesis of other vitamins was scarcer: 12 isolates for thiamine synthesis (belonging to *Bacillus, Halomonas, Qipengyuania, Roseibium, Oceanicaulis, Aquicoccus, Roseovarius*); four isolates for pyridoxal (vitamin B_6_) synthesis (belonging to *Halomonas* and *Marinobacter*); five for pimeloyl-ACP and biotin synthesis, including synthesis of the biotin precursor pimeloyl-ACP (belonging to *Bacillus, Halomonas, Marinobacter*). Only MMSF_3323 (*Halomonas* sp.) had the potential to synthesize the seven B vitamins that were investigated, while MMSF_3353 (*Bacillus* sp.) and all three *Marinobacter* isolates could synthesize five (Table S4). Provisioning of cobalamin by a separate *Halomonas* strain was demonstrated in the red alga *P. purpureum* (26). Therefore, there is high potential for B vitamin provisioning from Symbiodiniaceae-associated bacteria. How these nutrients are exchanged between the partners remains unknown, although some isolates did possess ABC transporters for thiamine that may facilitate its export. Intracellular bacteria may be able to directly export the vitamins in the Symbiodiniaceae cell’s cytoplasm, making it readily available.

The potential for bacterial B vitamin provisioning is important for several reasons. First, as shown in several other microalgae, B vitamin supplementation, specifically cobalamin, can increase algal growth (26, 30, 31). Second, cobalamin supplementation by bacteria was linked to increased thermal tolerance in *C. reinhardtii* (30), which is relevant to Symbiodiniaceae in the context of thermal stress and coral bleaching. Third, coral hosts of Symbiodiniaceae cannot synthesize B vitamins either. Some coral-associated bacteria, such as *Endozoicomonas*, have been hypothesized to synthesize B vitamins for their coral hosts (32–34). Symbiodiniaceae-associated bacteria may also provide B vitamins to the wider coral holobiont, rather than just Symbiodiniaceae.

### Potential for reactive oxygen species scavenging during thermal stress

ROS have been implicated in the thermal stress response of Symbiodiniaceae and in turn in coral bleaching (8, 9, 35). Because of this, the production of ROS is a key trait in the design of probiotics for improving thermal tolerance in Symbiodiniaceae and corals (18, 20, 36–39). While Symbiodiniaceae continuously produce ROS (such as singlet oxygen (^1^O_2_), superoxide (O_2_^−^), hydrogen peroxide (H_2_O_2_) and hydroxyl radicals (OH^-^)), thermal stress results in an increased production of ROS that may ultimately leak into coral cells and trigger coral bleaching (9). Thus, we looked at pathways and proteins potentially involved in ROS scavenging and/or the production of antioxidants. Pathways for the synthesis of glutathione and ubiquinone, two antioxidants, were complete in 30 and 4 isolates, respectively (Figure 2, Table S3). All Proteobacteria were able to produce glutathione, while only the four Gammaproteobacteria isolates (*Halomonas* and *Marinobacter*) were able to produce ubiquinone. Additionally, specific protein families (Pfam) or their synthesis pathways related to ROS and RNS scavenging were manually searched in the 49 genomes (Table 1): synthesis of zeaxhanthin, a carotenoid that quenches singlet oxygen and can scavenge other ROS (40); dimethylsulfonioproprionate (DMSP) and dimethylsulfate (DMS), OH^-^ scavengers (41); mannitol, another OH^-^ scavenger (42); ROS-scavenging enzymes, including superoxide dismutases (O_2_^-^ scavengers), peroxiredoxins and peroxidases (H_2_O_2_ scavengers), glutaredoxins and thioredoxins (general ROS scavengers). These are summarized in Figure 3.

**Figure 3:**
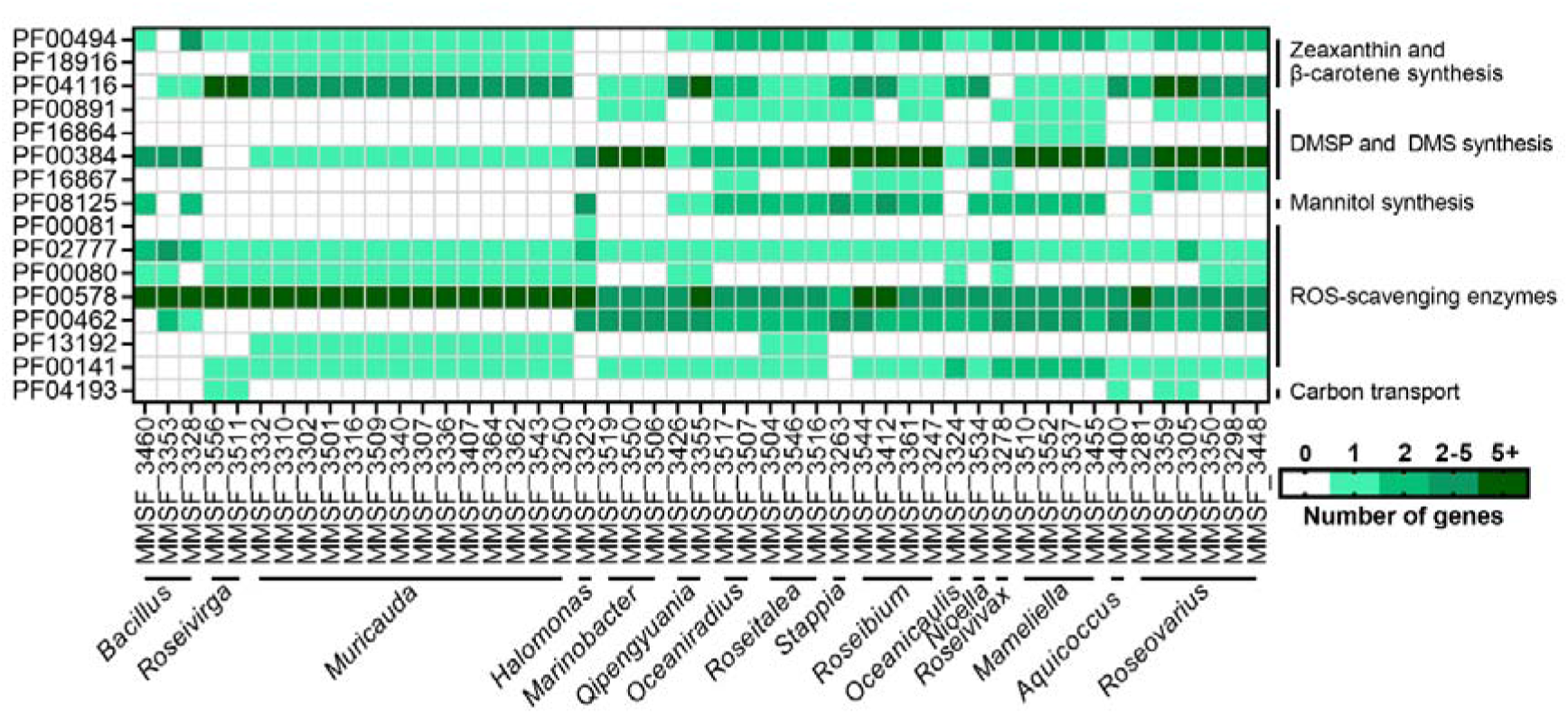
Putatively beneficial functions in Symbiodiniaceae-associated bacterial genomes. Protein families (Pfam) IDs were retrieved in each genome based on an InterProScan annotation. See Table 1 for full details on each Pfam ID. DMSP: dimethylsulfonioproprionate; DMS: dimethylsulfate; ROS: reactive oxygen species.

**Table 1:** Genes and respective protein families (Pfam) involved for which Symbiodiniaceae-associated bacterial genomes were screened. DMSP: dimethylsulfonioproprionate; DMS: dimethylsulfate; DMSO: dimethylsulfoxide; ROS: reactive oxygen species; SWEET: sugars will eventually be exported transporter.

| Pfam ID | Protein | Process |
| --- | --- | --- |
| PF00494 | Phytoene synthase | Zeaxanthin and β-carotene synthesis |
| PF18916 | Lycopene β-cyclase | |
| PF04116 | β-carotene hydroxylase | |
| PF00891 | Methyltransferase (DsyB) | DMSP and DMS synthesis |
| PF16864 | Methyltransferase (DsyB) |  |
| PF00384 | DMSO reductase |  |
| PF16867 | DMSP lyase |  |
| PF08125 | Mannitol 2-dehydrogenase | Mannitol synthesis |
| PF00081 | Superoxide dismutase [Mn] | ROS scavenging |
| PF02777 | Superoxide dismutase [Mn] |  |
| PF00080 | Superoxide dismutase [Cu-Zn] |  |
| PF00578 | Peroxiredoxin |  |
| PF00462 | Glutaredoxin |  |
| PF13192 | Thioredoxin |  |
| PF00141 | Peroxidase |  |
| PF04193 | SemiSWEET protein | Carbon transport |

Only 14 isolates, all belonging to *Muricauda*, were found to be able to synthesize zeaxanthin. This is consistent with a previous study showing *Durusdinium*-associated *Muricauda* can produce zeaxanthin and contributes to Symbiodiniaceae tolerance to thermal and light stress (20). Twenty-two isolates possessed the *dsyB* gene, involved in the production of DMSP (belonging to *Roseovarius, Mameliella, Roseovivax, Roseibium, Stappia, Roseitalea, Oceaniradius,* and *Marinobacter*), while all isolates except the two *Roseivirga* isolates were able to synthesize DMS, through DMSO reduction and/or DMSP cleavage. Interestingly, a *Roseovarius* isolate, MMSF_3448, which enhanced thermal tolerance in *Breviolum minutum* cultures (18) had the genes to produce both DMSP and DMS, which may be part of the mechanism underlying the benefit found in that study. Additionally, bacterial DMSP and DMS metabolism was shown to increase in the proximity of Symbiodiniaceae cells (43), suggesting sulfur-based interactions between Symbiodiniaceae and their associated bacteria. A gene encoding for a mannitol-1-phosphate dehydrogenase, necessary for mannitol production, was found in 22 isolates (belonging to *Bacillus, Halomonas, Qipengyuania, Oceaniradius, Aquicoccus, Mameliella, Roseivivax, Nioella, Roseibium, Stappia,* and *Roseitalea*). Mannitol is has been shown to mitigate thermal bleaching in corals and sea anemones (44, 45). Finally, while all isolates possessed ROS-scavenging enzymes, some isolates had particularly high numbers, including isolates belonging to *Halomonas*, *Muricauda, Roseibium*, and *Roseivirga*. Isolates of the *Muricauda* and *Roseitalea* genera were the only bacteria possessing genes encoding thioredoxins, while only *Roseitalea* isolates possessed all six queried enzymes. Hence, there is a high potential for Symbiodiniaceae-associated bacteria to assist with ROS removal during thermal stress via various mechanisms. This is particularly important for intracellular bacteria, as they may be able to export ROS-scavenging compounds directly into the Symbiodiniaceae cytoplasm, strengthen the antioxidant response, and minimize ROS leakage out of the Symbiodiniaceae cells. Most of the bacterial genera studied here are commonly detected in and cultured from cnidarian hosts (11, 46, 47) and may therefore participate in the *in hospite* Symbiodiniaceae response to thermal stress, and thereby influence the outcome of coral thermal stress. Nonetheless, whether these bacteria truly associate with Symbiodiniaceae *in hospite* remains unknown.

### Potential for carbon exchange

Finally, we investigated the presence of carbon exporters in the bacterial genomes. The “sugars will eventually be exported transporters” (SWEET) are bidirectional sugar transporters found in eukaryotes, and are hypothesized to regulate carbon exchange between Symbiodiniaceae and corals (48). SemiSWEET proteins are the bacterial homologues of SWEET proteins, although sugar export has not yet been confirmed in bacteria (49). Five isolates were found to have SemiSWEET genes (Figure 3): two *Roseivirga* isolates, one *Aquicoccus* isolate, and two *Roseovarius* isolates. In a healthy state, it is unlikely that bacteria would provide glucose to Symbiodiniaceae, which can make its own through photosynthesis, and the bacterial transporters may instead be used to import photosynthate. However, sugars may be exported via the same transporter, which may help compensate for the decrease in photosynthesis, and thereby glucose production, by Symbiodiniaceae during thermal stress. This is the first report of genes encoding for SemiSWEET proteins in Symbiodiniaceae-associated bacteria, and their functional analysis is needed to shed light on their role towards Symbiodiniaceae functioning and thermal tolerance.

### Genomic evolution in response to Symbiodiniaceae experimental evolution

Among the bacterial genomes sequenced from *C. proliferum* cultures, we obtained a representative from each of the three *C. proliferum* strains (WT10, SS03, and SS08) for four bacterial species, *Marinobacter* sp. (MMSF_3506, MMSF_3519, MMSF_3550), *Muricauda* sp. (MMSF_3501, MMSF_3509, MMSF_3543), *Mameliella* sp. (MMSF_3510, MMSF_3537, MMSF_3552), and *Roseitalea* sp. (MMSF_3504, MMSF_3516, MMSF_3546). Within a given bacterial species, all three genomes were >99.9% identical (based on ANI and AAI calculations, Figure S1), suggesting they belong to the same strain. Therefore, we assumed that the different bacterial isolates were present in the original *C. proliferum* culture, before experimental evolution started, and independently evolved in the three *C. proliferum* strains, under elevated temperature (SS strains) versus ambient temperature (WT10 strain). We performed a single nucleotide polymorphism (SNP) analysis to uncover which bacterial genes, if any, were impacted by host heat selection.

Across the four bacterial strains, we detected 32 SNPs between bacteria isolated from WT10 *C. proliferum* and the corresponding bacteria isolated from SS *C. proliferum* (Table S5). Six SNPs were synonymous mutations and therefore not expected to bear phenotypic effects. For each given bacterial strain, we next chose to focus on the non-synonymous mutations that were present in both bacteria isolated from SS *C. proliferum* (Table S5, boldened rows), as there is a higher chance these would be true SNPs and may point to convergent evolution under heat selection. No non-synonymous SNPs were detected in *Roseitalea* sp.

In *Mameliella* sp., a missense mutation affected the *chvI* gene, a transcriptional regulator known to be essential for the virulence of the plan pathogen *Agrobacterium tumefaciens* (50), as well as successful establishment of symbiosis in the nodule-forming and nitrogen-fixing plant associates *Rhizobium leguminosarum* and *Sinorhizobium meliloti* (51, 52).

In *Muricauda* sp., missense mutations affected genes encoding a heme A synthase, a histidine kinase, and a DNA gyrase. Histidine kinases have a broad range of actions in signal transduction, while DNA gyrases are involved in DNA replication. The impact of mutations on those two genes is therefore hard to assess. Heme A synthases are involved in heme synthesis, a cofactor involved in many cellular processes, including antioxidant functions through its incorporation into hemoproteins, as well as infectivity and virulence. Intracellular *Brucella abortus* are dependent on heme production for their survival (53), while heme deficiency negatively impacts mouse tissue colonization by *Staphylococcus aureus* (54). Additionally, excess free heme catalyzes the formation of ROS, resulting in oxidative stress (55). If this mutation resulted in decreased heme synthesis, it may contribute to lower ROS production during heat stress and thereby participate in the heat tolerance of SS *C. proliferum*.

In *Marinobacter* sp., a missense mutation affected *gspE*, which encodes an ATPase involved in the T2SS. As is the case with most secretion systems, the T2SS is required for the virulence of a variety of pathogens (56), as well as the colonization of leech gut by the mutualistic symbiont *Aeromonas veronii* (57). Additionally, a frameshift mutation affected *pilO*, while a mutation in *pilM* resulted in the loss of a stop codon; both genes are involved in the type IV pilus assembly. Type IV pili are involved in the formation of biofilms, but also support bacterial motility and may play a role in host colonization (58).

In all three cases, the detected mutations in response to heat selection affected genes potentially involved in host-symbiont interactions, such as host infection or symbiosis establishment. Whether they enhance or are detrimental to these interactions remains unknown, and functional studies will be needed to uncover the phenotypic effects of these mutations.

## Conclusion

We provide the first extensive genomic analysis of Symbiodiniaceae-associated bacteria, by presenting the genomes of 49 bacteria isolated from six Symbiodiniaceae genera. Most bacteria have the potential to be beneficial for their hosts, through the production of B vitamins or antioxidants. *Halomonas* sp. MMSF_3323 is particularly interesting as it was predicted to synthesize seven essential B vitamins. Isolates of the *Muricauda* genus may also be crucial as they were the only bacteria in our data set that are able to produce antioxidant carotenoids. *Marinobacter* sp. are also likely important for Symbiodiniaceae health as they can produce four B vitamins, as well as many ROS-scavenging compounds, such as ubiquinone and glutathione. It is noteworthy that the functional potential of these bacteria was independent of their Symbiodiniaceae host. Given that these Symbiodiniaceae were initially obtained from corals, clams, and anemones, it is unclear whether the bacteria analyzed here would interact directly with an animal host *in hospite*, or rather only with Symbiodiniaceae. Genomic analyses of Symbiodiniaceae-associated bacteria *in hospite* are therefore needed to assess potential lifestyle differences.

Finally, we analyzed the effect of heat selection on genome evolution in four bacterial species. In *Muricauda* sp., *Marinobacter* sp., and *Mameliella* sp., mutations observed in bacteria isolated from heat-evolved Symbiodiniaceae were observed in genes potentially involved in host-symbiont interactions. While the effect of the mutations was not investigated, it is possible that future climate conditions will reinforce or damage interactions between bacteria and Symbiodiniaceae. Whether this may affect coral climate resilience remains to be investigated. More extensive genomic comparisons, using long-read sequencing for increased accuracy, are needed to fully understand the response of Symbiodiniaceae-associated bacteria to host heat selection. Overall, our study reveals the beneficial potential of Symbiodiniaceae-associated bacteria.

## Material and methods

### Cultures of bacteria from three *Cladocopium proliferum* strains

Three *Cladocopium proliferum* cultures were selected to isolate pure bacteria from: SCF055-01.10 (WT10), SCF055-01.03 (SS03), and SCF055-01.08 (SS08). These three *C. proliferum* strains originated from a monoclonal culture isolated from *Acropora tenuis* (59). SS03 and SS08 have been evolving in the laboratory after ratcheting to 31°C since 2011 while WT10 was kept at 27°C. At the time of sampling for this study (August - October 2022), cultures were experimentally evolved for an estimated 300-330 generations (based on generation time of the WT cultures). Symbiodiniaceae cultures were maintained in 15 mL Daigo’s IMK medium (1X), prepared with filtered Red Sea Salt water (fRSSW, 34 ppt) in sterile 50 mL polypropylene culture flasks where media was changed fortnightly. These flasks were kept in a 12 h light:12 h dark incubator with lighting at 50–60 μmol photons m^−2^ s^−1^ of photosynthetically active radiation (Taiwan Hipoint Corporation, model 740FHC LED, light chambers) and temperature at either 27°C (WT) or 31°C (SS).

Symbiodiniaceae cultures cell counts were quantified (Life Technologies Countess II FL) and an aliquot of 10^6^ cells was deposited in a 5-μM mesh size strainer (pluriSelect, Germany) to separate the Symbiodiniaceae (ranging from 6-9 μm in diameter) and their closely associated (physically attached/intracellular) bacterial cells from loosely associated (planktonic) bacterial cells. Strainers were sealed with parafilm and centrifuged at 10,000 × *g* for 3 min. A volume of 500 μL of fRSS was added to the strainers, which were sealed and centrifuged at 10,000 × *g* for 3 min. Symbiodiniaceae cells and their associated bacteria were resuspended from the filter in 1 mL of fRSS. For each culture, one aliquot was not processed any further and one other aliquot was bead beaten with 100 mg of sterile glass beads (400-600 nm) for 30 s at 30 Hz in a Tissue-lyser II (Qiagen, Germany), in order to open up Symbiodiniaceae cells and release intracellular bacteria to facilitate their growth on agar plates. Both aliquots were serially diluted spread onto BD Difco™ Marine Agar 2216 (MA) and three Oxoid R2A Agar (R2A) prepared with fRSS, culture plates. Plates were incubated at 27°C (for the WT10 samples) or at 31°C (SS03 and SS08 samples) for 7 d to facilitate bacterial colony growth. Representatives of all morphologies were subcultured to attain culture purity. Pure cultures were stored in sterile 40% glycerol at −80°C.

### 16S rRNA gene sequencing from cultured bacteria

Individual pure freshly grown bacterial colonies were suspended in 20 μL sterile Milli-Q® water, incubated for 10 min at 95°C then used as templates in PCRs. PCR amplification of the bacterial 16S rRNA gene was performed with primers 27F and 1492R (60) in reactions containing 2 μL template DNA, 1 μL of each primer at a 10 μM concentration (0.25 μM final concentration), 20 μL of 2X Mango Mix (Bioline 25034) and 16 μL of PCR grade water. The amplification cycle was: 5 min at 94°C; 30 cycles of 60 s at 94°C, 45 s at 50°C, and 90 s at 72°C; 10 min at 72°C; with a final holding temperature of 4°C. The PCR products were purified Sanger sequenced with primer 1492R at Macrogen (South Korea). Raw sequences were trimmed and proofread in Geneious Prime v2019.1.3. Sequences were analyzed in BLAST. For each sequence, the closest hit and percent identity was recorded in Table S1.

### DNA extraction and whole-genome sequencing

For each isolate selected for whole-genome sequencing, a single colony was picked with an inoculation loop and snap-frozen. DNA extraction was performed on the QIAsymphony using the DSP Virus/Pathogen Mini Kit (Qiagen). Library preparation performed using Nextera XT (Illumina Inc.) according to manufacturer’s instructions. Whole-genome sequencing was performed on NextSeq 500/550 with a 150bp PE kit.

### Genome assembly and annotation

Raw reads were trimmed and quality-filtered using Trimmomatic v0.36 (61) (HEADCROP:10 LEADING:5 TRAILING:5 SLIDINGWINDOW:4:28 MINLEN:30). The quality of raw reads before and after trimming was checked with FASTQC v0.11.9 (62). Trimmed and quality-filtered reads were de novo assembled into contigs using SPAdes v3.15.5 (63) with 21, 33, 55 and 77 k-mers and the option “--careful” was applied to minimise the number of mismatches and short indels. All contigs with a length of <1000 bp were removed using BBMap v38.96 (64). Subsequently, the levels of completeness and contamination of assembled genomes were assessed using CheckM v1.2.2 using the “lineage_wf” workflow (65). Taxonomic assignment of all assembled genomes was carried out using GTDB-Tk v2.1.0 (66) using the “classify_wf” workflow. GTDB-Tk assigned the taxonomy of the genomes based on 120 bacterial marker genes. A phylogenetic tree was built in IQ-Tree v2.0.3 (67) using the best model LG+F+R4, selected by ModelFinder wrapped in IQ-tree (68), and 1000 ultrafast bootstrap replicates (69). The tree was visualized in iTOL v6 (70). Average Nucleotide Identities (ANI) and Average Amino acid Identities (AAI) of all assembled genomes were calculated using a genome-based matrix calculator (71) and plotted in R using the ggplot2 package (72).

Gene prediction was performed in Bakta v1.7.0 (73) and InterProScan v5.55 using with the parameter “-appl pfam” and evalue <1e-5 (74). Metabolic pathways, transport systems and secretion systems were annotated via METABOLIC-G v4.0 (75) applying “-m-cutoff 0.75” to include pathways which are ≥75% complete. The completeness of metabolic pathways (KEGG module database, https://www.genome.jp/kegg/module.html), transporters and secretion systems was estimated in EnrichM v0.6.4 (76). KEGG modules of interest were then plotted in GraphPad Prism. Pfam IDs of interest (Table 1) were queried in the InterProScan annotation, and the number of hits for each Pfam IDs per genome was plotted in GraphPad Prism.

### Single nucleotide polymorphism analysis

Sequencing reads were assembled using the genome assembler pipeline Shovill v1.1.0 (77). Briefly, the Shovill pipeline included read trimming using Trimmomatic v0.39 (61), de novo assembly with SPAdes v3.15.5 (63) and genome polishing with Pilon v1.24 (78). After the pipeline, additional polishing was performed by mapping the reads back to the contigs with BWA v0.7.17 (79) and sorting the resulting SAM/BAM files using SAMtools v1.15.1 (80). Pilon v1.24 was then used to correct bases, fix mis-assemblies and fill gaps. The reformat.sh script from the Bbmap package v38.76 (-minlength=1000) was used to filter out contigs less than 1000bp (<u>sourceforge.net/projects/bbmap/</u>). The quality and completeness of the assemblies was assessed with CheckM v1.2.1 (65), BUSCO v5.3.2, and CAT/BAT v5.2.3. The draft genome assemblies were then annotated with Bakta v1.7.0 (73).

Single nucleotide polymorphism (SNP) detection between WT and SS samples were then performed using snippy v4.6.0 (81), where both WT and SS samples were inputted as the reference genome in turn. Variants were then selected if they were found in both the reciprocal snippy analyses.

## Data availability

Genbank accession numbers for 16S rRNA sequences of cultured bacteria are OQ945078-OQ945142 (see Table S1). Genome assemblies and raw NextSeq reads are available under NCBI BioProject ID PRJNA971760 (see Table S2 for individual accession numbers). A subset of the snippy output files is available on Zenodo at 10.5281/zenodo.8210291.

## Acknowledgements

This research was supported by the Australian Research Council (FL180100036 to MJHvO) and the Native Australian Animals Trust (to JM). JL was supported by a Future Fulbright scholarship. We thank Glen Carter for assistance with genome sequencing; Kshitij Tandon and Torsten Seemann for assistance with genome analyses; the Melbourne Research Cloud, the University of Melbourne’s Research Computing Services and the Petascale Campus Initiative for providing the high-performance computing instances needed for this work.

## Author contributions

Conceptualization: JM, LLB, MJHvO; Formal analysis: JM, GKP; Funding acquisition: JM, MJHvO; Investigation: JM, JL, LMJ; Methodology: JM, JL; Visualization: JM, GKP; Writing - Original Draft: JM, MJHvO. All authors read and edited the manuscript.

