## Supplementary material for "Functional potential and evolutionary response to long-term heat selection of bacterial associates of coral photosymbionts": Figure S1

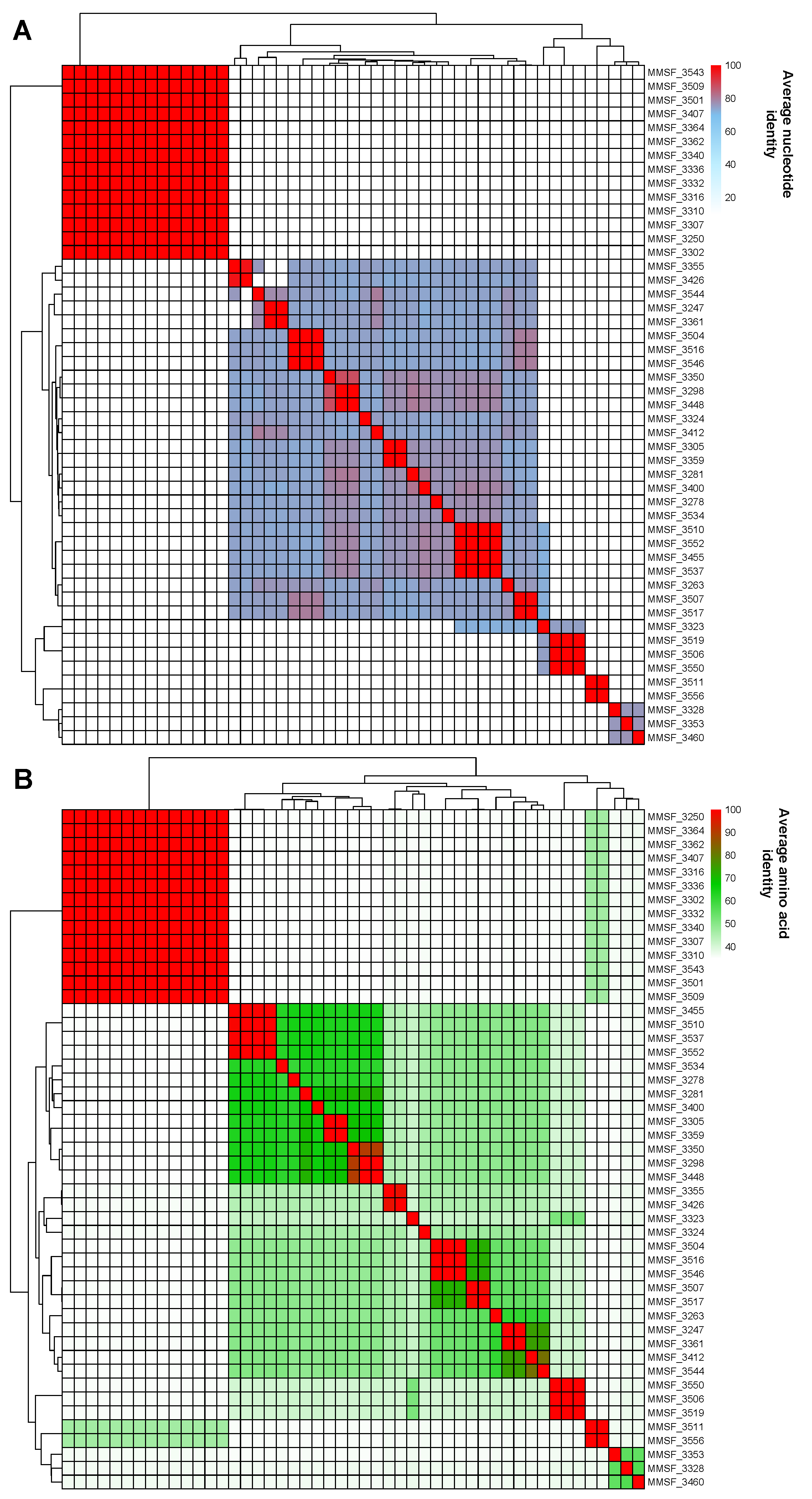

**Figure S1:** Average nucleotide identity (ANI) (A) and average amino acid identity (AAI) (B) of 49 genomes of Symbiodiniaceae-associated bacteria.

**Table S1:** Database of bacteria isolated from three *Cladocopium proliferum* strains. For each isolate: source Symbiodiniaceae strain, closest BLAST result and full taxonomy are provided. (attached)

**Table S2:** Characteristics of assembled bacterial genomes isolated from Symbiodiniaceae cultures, including Symbiodiniaceae host characteristics, genome size, taxonomy, completeness, and contamination. (attached)

**Table S3:** Metabolic pathway annotation and completeness of bacterial genomes isolated from Symbiodiniaceae. Pathways were annotated with METABOLIC-G and completeness was estimated using EnrichM. (attached)

**Table S4:** Number of isolates in each genus with complete (>75%) pathways for B vitamin biosynthesis. Cells are empty when no isolate has a complete pathway. (attached)

**Table S5:** Single nucleotide polymorphisms (SNPs) detected between genomes belonging to the same bacterial strain (either *Mameliella* sp., *Muricauda* sp., *Marinobacter* sp., or *Roseitalea* sp.). Within each bacterial strain, one bacterial genome obtained from a WT10 *C. proliferum* culture was compared to two bacterial genomes obtained from SS03 and SS08 *C. proliferum* cultures (WT vs SS comparisons). For each comparison, the reciprocal (SS vs WT) was also performed, and only SNPs detected in both comparisons were considered true SNPs. Within each bacterial strain, non-synonymous mutations detected in both genomes isolated from SS *C. proliferum* are highlighted in bold. (attached)
